# Peripherally expressed misfolded proteins remotely disrupt brain function and aggravate stroke-induced brain injury

**DOI:** 10.1101/785477

**Authors:** Yanying Liu, Kalpana Subedi, Aravind Baride, Svetlana Romanova, Christa C. Huber, Xuejun Wang, Hongmin Wang

## Abstract

Impaired proteostasis has been linked to various diseases, whereas little is known about the impact of peripherally misfolded proteins on the brain. We here studied the brain of mice with cardiomyocyte-restricted overexpression of a missense (R120G) mutant small heat shock protein, αB-crystallin (CryAB^R120G^). At baseline, the CryAB^R120G^ mice showed impaired cognitive and motor functions, aberrant protein aggregates, neuroinflammation, impaired blood-brain barrier, and reduced proteasome activity in the brain compared with their non-transgenic (Ntg) littermates. Ischemic stroke dramatically exacerbated these pathological alterations and caused more severe brain dysfunction in CryAB^R120G^ mice than in the Ntg mice. Intravenously injecting the exosomes isolated from CryAB^R120G^ mouse blood into wild-type mice caused the similar phenotypes seen from CryAB^R120G^ mice. Importantly, the CryAB^R120G^ protein showed the prion-like properties. These results suggest that peripherally misfolded proteins in the heart remotely disrupt brain function through prion-like neuropathology, which may represent an underappreciated mechanism underlying heart-brain crosstalk.

---

Maintaining the integrity of the proteome is essential for cell survival and normal functioning. However, proteins frequently misfold due to genetic mutations, stress conditions, or unique metabolic challenge conditions^1^. Mammalian cells evolve three major mechanisms to maintain protein homeostasis, or proteostasis, i.e. molecular chaperones, the ubiquitin-proteasome system (UPS) and the autophagy lysosome system^2^. These cellular surveillance mechanisms detect and repair misfolded proteins or, in many situations, completely annihilate the misfolded proteins from the inside of cells^3^. Conversely, impaired protein quality control causes proteotoxicity that has been linked to numerous devastating human diseases known as protein conformational disorders, including many neurodegenerative diseases, metabolic disorders, cardiomyopathies, liver diseases and systemic amyloidosis^3^.

Data from clinical studies suggest a close link between heart disorders and neurological deficits, although the mechanism underlying the connection remains unclear. The neurological deficits associated with heart diseases are generally recognized as a common adverse consequence of reduction in cerebral blood supply, alterations of cerebrovascular reactivity, and modification of blood pressure^4, 5^. Apart from these physiological alterations, few studies examine whether some specific circulating factors associated with a heart disease can directly modulate brain function, and whether these factors influence the integrity of the brain-blood barrier (BBB) and the outcomes following acute brain injury, such as ischemic stroke.

Stroke is the fifth most common cause of death and is a leading cause of serious long-term disability in the United States^6^. Ischemic stroke is caused by blockage of cerebral artery and is associated with neuronal loss and dysfunction. Following cerebral ischemia and reperfusion (I/R), mitochondria dysfunction, glutamate-induced excitotoxicity, and neuroinflammation occurs, leading to oxidative stress, protein damage and aggregation^7, 8^. Proteostasis has been shown to play an important role in neuronal injury and functional recovery following I/R^9, 10^. However, very little is known about the impact of peripherally impaired proteostasis on ischemic stroke-induced brain injury and functional recovery.

Unlike stroke, many other neurological diseases involve specific misfolded pathogenic proteins that aggregate into seeds that serve as self-propagation agents for the progression of diseases^11^. This prion-like property of pathogenic proteins has been seen in the pathology of Lewy body^12, 13^ and α-synuclein^14, 15^, β-amyloid^16, 17^, mutant huntingtin^18^, and various other neurodegenerative disease-associated proteins^19^. These studies focused on cell-cell pathological propagation between grafted cells and host cells in the brain. However, it remains unknown whether such prion-like phenomenon can be propagated from peripheral tissues to the brain, leading to functional or pathological alterations in the brain or exacerbation of I/R induced brain injury. To address this question, we here investigated the brain of a transgenic mouse expressing a missense (R120G) mutant alphaB-crystallin (CryAB^R120G^), selectively in the cardiomyocytes, which causes desmin-related cardiomyopathy^20, 21^. CryAB, also known as HSPB5 is a small heat shock protein that is expressed at high levels in the lens but is also ubiquitously expressed in other tissues such as the heart and skeletal muscles^22^. A majority of this line of CryAB^R120G^ mutant mice died of congestive heart failure between 6 and 7 months (m) of age, with pronounced accumulation of desmin and CryAB aggregates and impaired UPS function in the heart, indicating impaired proteostasis in the cardiomyocytes^20^. This allows us to assess whether the animal during the cardiac compensatory stage exhibit any functional or pathological alterations in the brain prior to death in the absence or presence of transient I/R insult.

## Results

### Mice with cardiomyocyte-restricted expression of the CryAB^R120G^ protein showed impaired cognitive behaviors and reduced motor functions

The heart function of the CryAB^R120G^ mice is known at the compensatory stage at 3 months (m) of age, showing increased contractile function and reduced relaxation function.^20^ At this age, our behavioral tests revealed reduced spatial learning and memory capability compared their non-transgenic (Ntg) littermates, as demonstrated in the radial arm water maze (RAWM) test (Fig. 1a), Y-maze test (Fig. 1b), and novel object recognition test (Fig. 1c). In the open field test at 3 m, the CryAB^R120G^ mice spent less time in the center of the field than the Ntg mice (Fig. 1d) and showed decreased exploratory behavior (Fig. 1e), suggesting increased anxiety and decreased motor function in the CryAB^R120G^ transgenic mice. The reduced motor function in CryAB^R120G^ mice was further supported by the string agility test at 3 m (Fig. 1f) and rotarod test at 4 m (Fig. 1g). These altered cognitive and locomotion behaviors persisted in the CryAB^R120G^ mice until they died around 6 m (data not shown), indicating that cardiomyocyte-restricted expression of the CryAB^R120G^ protein disrupts brain functions.

**Fig. 1.**
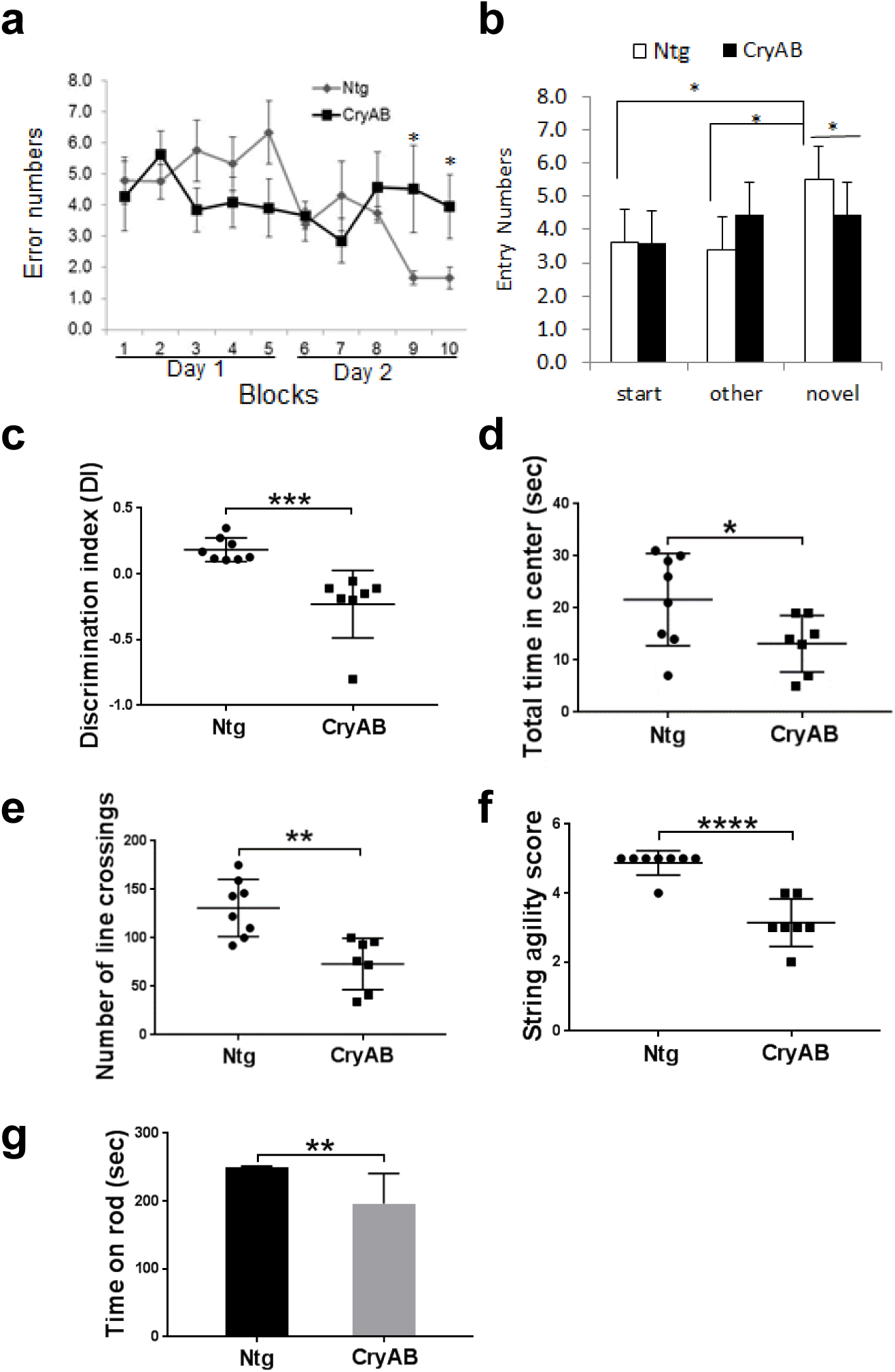
Mice with cardiomyocyte-restricted expression of CryAB^R120G^ showed impaired cognitive and motor functions. **a** Measurement of mouse spatial learning and memory in CryAB mice and their Ntg littermates by RAWM test at 3 m. **b** Y maze test of mice at the age of 3 m. **c** Object recognition test at 3 m. **d** and **e** Open field test at 3 m including the total time spend in center **d** and total lines crossing **e**. **f** String ability test of mice at 3 m. **g** Rotarod test at 4 m. Data are shown as meanLJ±LJS.D.; n = 8 for Ntg mice and n = 7 for CryAB mice. \**p* < 0.05, \*\**p* < 0.01, \*\*\**p* < 0.001, \*\*\*\**p* < 0.0001.

### Mice with cardiomyocyte-restricted expression of the mutant CryAB^R120G^ protein showed protein aggregates in the brain

As previous data have shown that the CryAB^R120G^ mice show pronounced accumulation of desmin and CryAB aggregates and impaired UPS function in the heart^20^, we next examined whether the cardiomyocyte-restricted expression of the misfolded CryAB^R120G^ protein is associated with any pathological alterations in the brain. Accordingly, following completion of the behavioral tests, the mice were euthanized and their brains were used for various pathological analyses at 5 m. Thioflavin-S staining, a validated protein staining method to quantify amyloid protein aggregates^23^, showed a remarked increase of fluorescence intensity in the brain of CryAB^R120G^ mice compared to Ntg mice (Fig. 2a, b). Immunostaining of brain sections with a CryAB specific antibody showed increased immuno-reactivity (Figs. 2c, d) and CryAB aggregates in the cells of CryAB^R120G^ mouse brains (Fig. 2c). The aggregates in CryAB mouse brains were mainly located in microglia and neurons, as CryAB was colocalized with a microglia-specific marker, Iba1 (Fig. 2e), and a neuron-specific marker, NeuN (Fig. 2f). Further, more Iba1-positive cells were seen in the brains of CryAB^R120G^ mice than those in Ntg mice (Fig. 2g, h), indicative of inflammation in the brain. In further support of this, the CryAB^R120G^ brains also exhibited upregulation of two immunoproteasome subunits, β1i and β5i in the cortex (Fig. 2i, j). Intriguingly, CryAB^R120G^ mouse brain displayed reduced proteasome activity (Fig. 2k) and increased accumulation of polyubiquitinated proteins (Fig. 2l, m) and K48-linked polyubiquitinated proteins (Fig. 2n, o) in the cortex. These results suggest that expression of the mutant CryAB^R120G^ protein in the heart impairs proteostasis in the brain, causing protein aggregation, neuroinflammation and proteasome dysfunction.

**Fig. 2.**
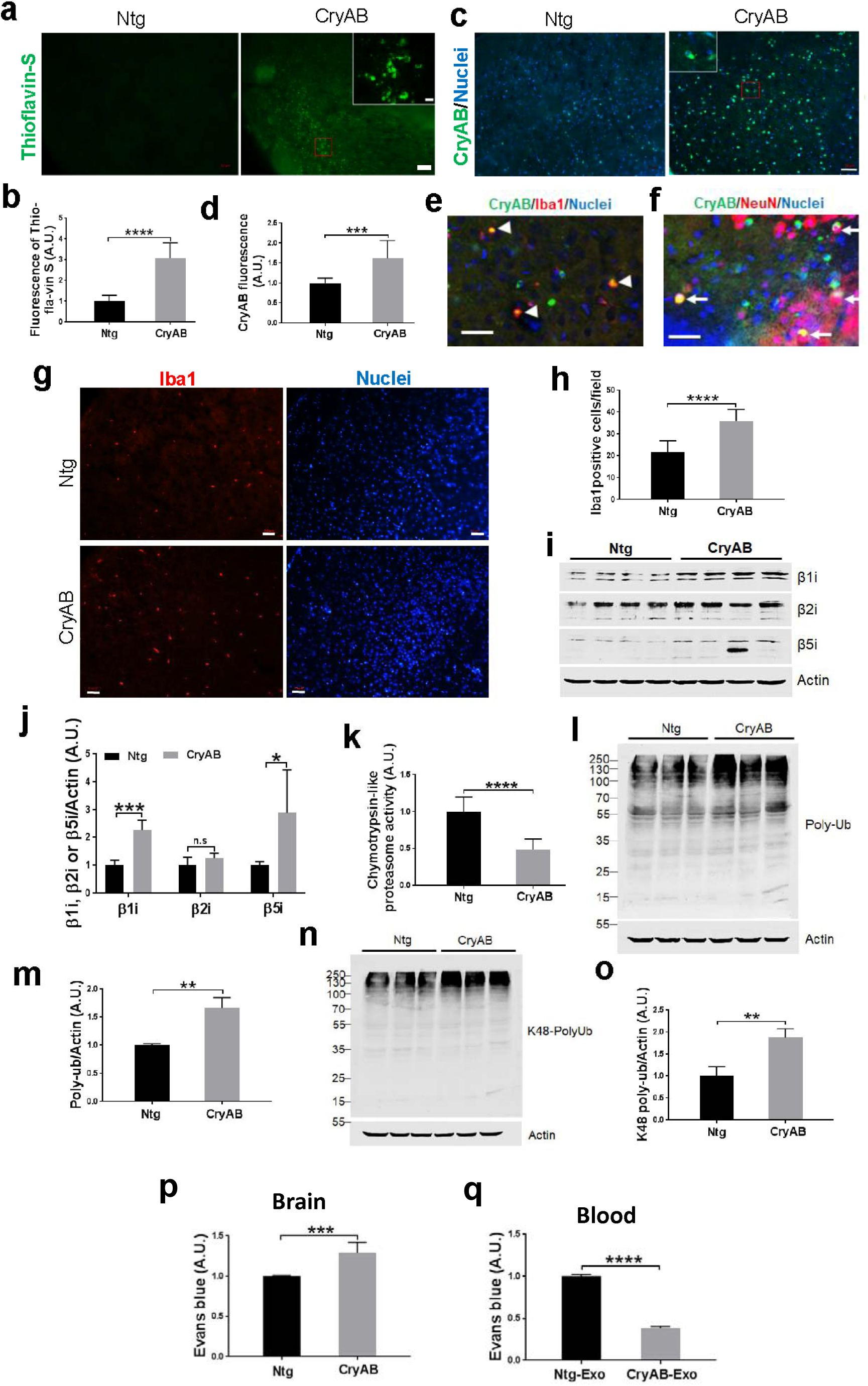
The brain of mice with cardiomyocyte-restricted expression of CryAB^R120G^ showed increased protein aggregates and neuroinflammation, and impaired proteasome functionality and BBB. **a** Thioflavin S staining of brains at age of 5m. Scale bars: 50 µm. **b** Quantitative analysis of **a**. **c** Immunohistochemistry analysis of CryAB immunoreactivity intensity (green) in the brain at 5m. Hoechst 33342 was used to stain cell nuclei. Scale bars, 50 μm. **d** quantitative analysis of **c**. **e** Immunostaining of both CryAB (green) and Iba1 (red) in the cortex of CryAB mouse brain. Arrowheads, colocalised cells. Scale bar, 50 μm. **f** Immunostaining of both CryAB (green) and NeuN (red) in the cortex of CryAB mouse brain. Arrows, colocalized cells. Scale bar, 50 μm. **g** Immunostaining of Iba1-postive cells in the Ntg or CryAB brain cortex. Red: Iba1, blue: Hoechst 33342. Scale bars, 50 μm. **h** Quantitative analysis of **g**. **i** Western blot analysis of immunoproteasomes expression in the brain cortex at 5m. **j** Quantitative analysis of **i**. **k** Proteasome activity in the cortex at 5m. **l** Western blot analysis of polyubiquitinated proteins in brain cortex at 5m. **m** Quantitation of **l**. **n** Western blot analysis of K48-linkage polyubiquitinated proteins in the brain cortex at 5m. **o** Quantitation of **n**. **p** Increased penetration of the low molecular weight BBB tracer, Evans blue, was observed in brain at 3 m of mice. **q** Evans blue concentration was reduced in the blood. All numerical data are shown as mean ± SD; n = 3 or 4. *\*p < 0.05,* \*\**p* < 0.01, \*\*\**p* < 0.001, \*\*\*\**p* < 0.0001.

To determine how cardiomyocyte-restricted expression of misfolded CryAB^R120G^ proteins causes cerebral neuropathological alterations, we next assessed whether the CryAB^R120G^ mouse is associated with impaired blood-brain barrier (BBB), as the BBB plays a key role in maintaining the normal function of the brain^24^. Accordingly, we intraperitoneally injected the Evans blue dye, an inert tracer that has been commonly used to measure vascular permeability in animal models^25^, into both CryAB^R120G^ and their non-transgenic (Ntg) littermate mice at 3 m, when cognitive and motor dysfunction occurred as described above. After 1 h, we observed a significantly higher level of the Evan blue in the CryAB^R120G^ mouse brain than in the Ntg mouse brain (Fig. 2p). This result was further supported by a reduced Evan blue level in CryAB^R120G^ mouse blood when compared to Ntg mouse blood (Fig. 2q), indicating that CryAB^R120G^ mice show increased permeability of BBB.

### Cardiomyocyte-restricted expression of misfolded CryAB^R120G^ exacerbated I/R-induced brain injury and cognitive and motor defects

To determine whether peripherally impaired proteostasis influences brain injury and functional recovery, we next challenged both CryAB^R120G^ male mice and their male Ntg littermates with I/R at 2-3 m and then examined brain injury and functional recovery. Following 45 minutes of MCAO and 24 hours of reperfusion (Fig. 3a), CryAB^R120G^ mice showed increased infarct volume compared to their Ntg littermates (Fig. 3b, c). This result was further confirmed by the Nissl staining data, which showed fewer survival neurons in CryAB^R120G^ mouse brains than Ntg mouse brains after 7 days of reperfusion (Fig. 3d, e). Consistently, CryAB^R120G^ mice also showed a higher mortality rate (Fig. 3f) and more severe neurological deficits (Fig. 3g) than their Ntg littermates. These observations were further supported by other behavioral tests following I/R. After 14 days of reperfusion, CryAB^R120G^ mice exhibited reduced motility compared with Ntg mice in the string agility (Fig. 3h) and the rotarod test (Fig. 3i). CryAB^R120G^ mice also showed poorer performance in the object recognition test (Fig. 3j), Y-maze test (Fig. 3k) and radial-arm water maze test (Fig. 3l) than their Ntg littermates, indicative of exacerbated cognitive dysfunctions in CryAB^R120G^ compared to Ntg mice. Thus, peripheral expression of misfolded CryAB^R120G^ protein aggravates I/R-induced brain injury and neurological deficits.

**Fig. 3.**
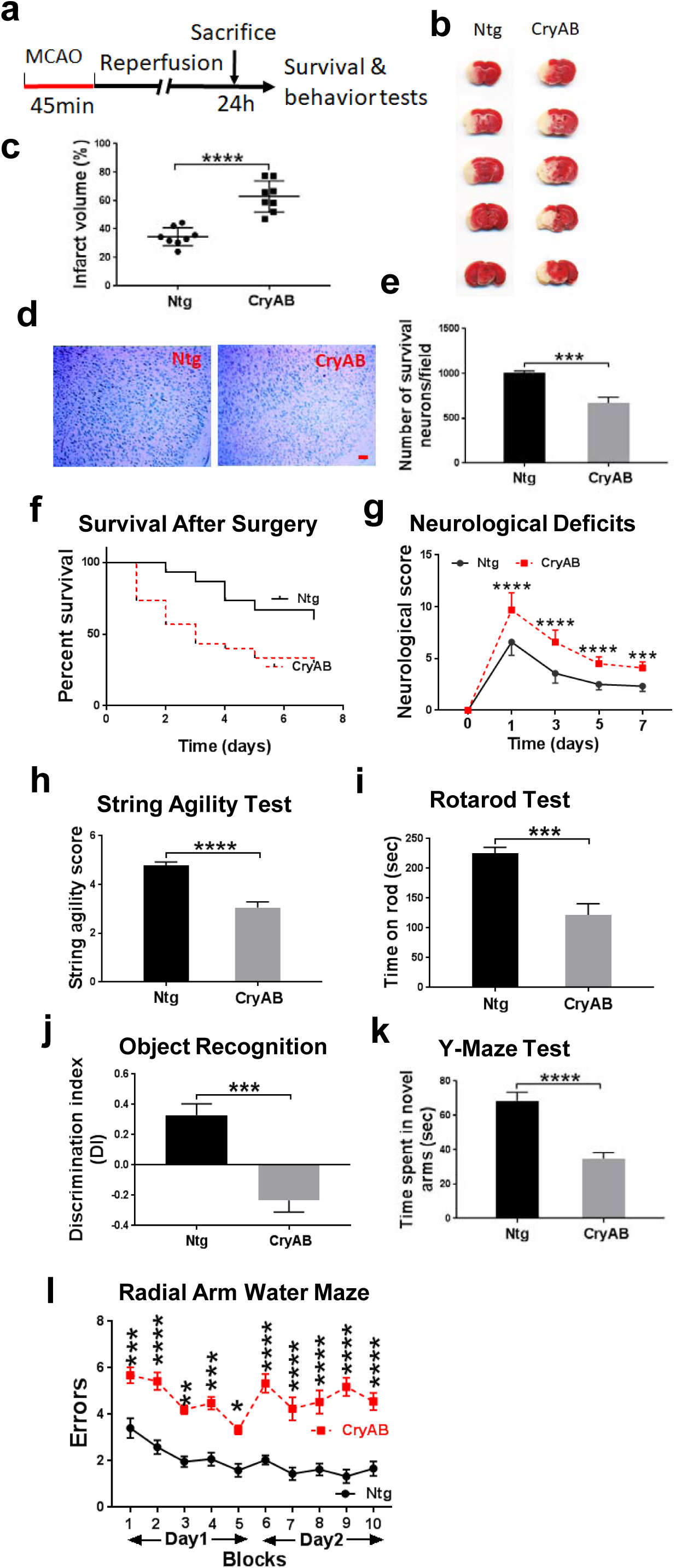
Cardiomyocyte-restricted expression of misfolded CryAB^R120G^ protein exacerbated I/R-induced brain injury and cognitive and motor deficits. **a** Diagram of the experimental design. **b** TTC staining mouse brains after 24 hours of reperfusion. **c** Quantitative analysis of **b**. **d** Representative images of Nissl staining results after 7 days of reperfusion. **e** Quantitative analysis of **d**. **f** Survival rate of mice after MCAO. **g** Functional recovery of mice after MCAO. **h** String agility test result after 14 days of reperfusion. **i** Rotarod test result after 14 days of reperfusion. **j** Object recognition test result after 14 days of reperfusion. **k** Y-maze test result after 14 days of reperfusion. **l** Radial arm water maze (RAWM) result after 14 days of reperfusion. Numerical data are shown as mean ± SEM; n = 9. * *p* < 0.05, ** *p* < 0.01, *** *p* < 0.001, **** *p* < 0.0001.

### Cardiomyocyte-restricted expression of misfolded CryAB^R120G^ exacerbated I/R induced neuroinflammation

To determine whether cardiomyocyte-restricted expression of misfolded CryAB^R120G^ proteins alters I/R triggered neuroinflammation, we immunohistochemically examined astrocytes and microglia in the mouse brains after 2 days of I/R. Both GFAP (glial fibrillary acidic protein, an astrocyte marker) (Fig. 4a, b) and Iba1 (a microglial marker) (Fig. 4c, d) showed a higher level of immunoreactivity in the CryAB^R120G^ mouse brains than in the Ntg brains, indicating increased activation of astrocytes and microglia in CryAB^R120G^ mouse brain following I/R. Importantly, the immunoreactivity of TNFα, a potent activator of the immune system, astrocytes, and microglia, was also higher in CryAB^R120G^ mouse brain than in Ntg mouse brain after 2 days of I/R (Fig. 4e, f), suggesting pronounced neuroinflammation occurring in CryAB^R120G^ mouse brains. This was further supported by the upregulation of all of the three immunoproreasome subunits, β1i, β2i and β5i (Fig. 4g, h). Collectively, these data reveal that cardiomyocyte-restricted expression of misfolded CryAB^R120G^ protein enhances I/R induced neuroinflammation.

**Fig. 4.**
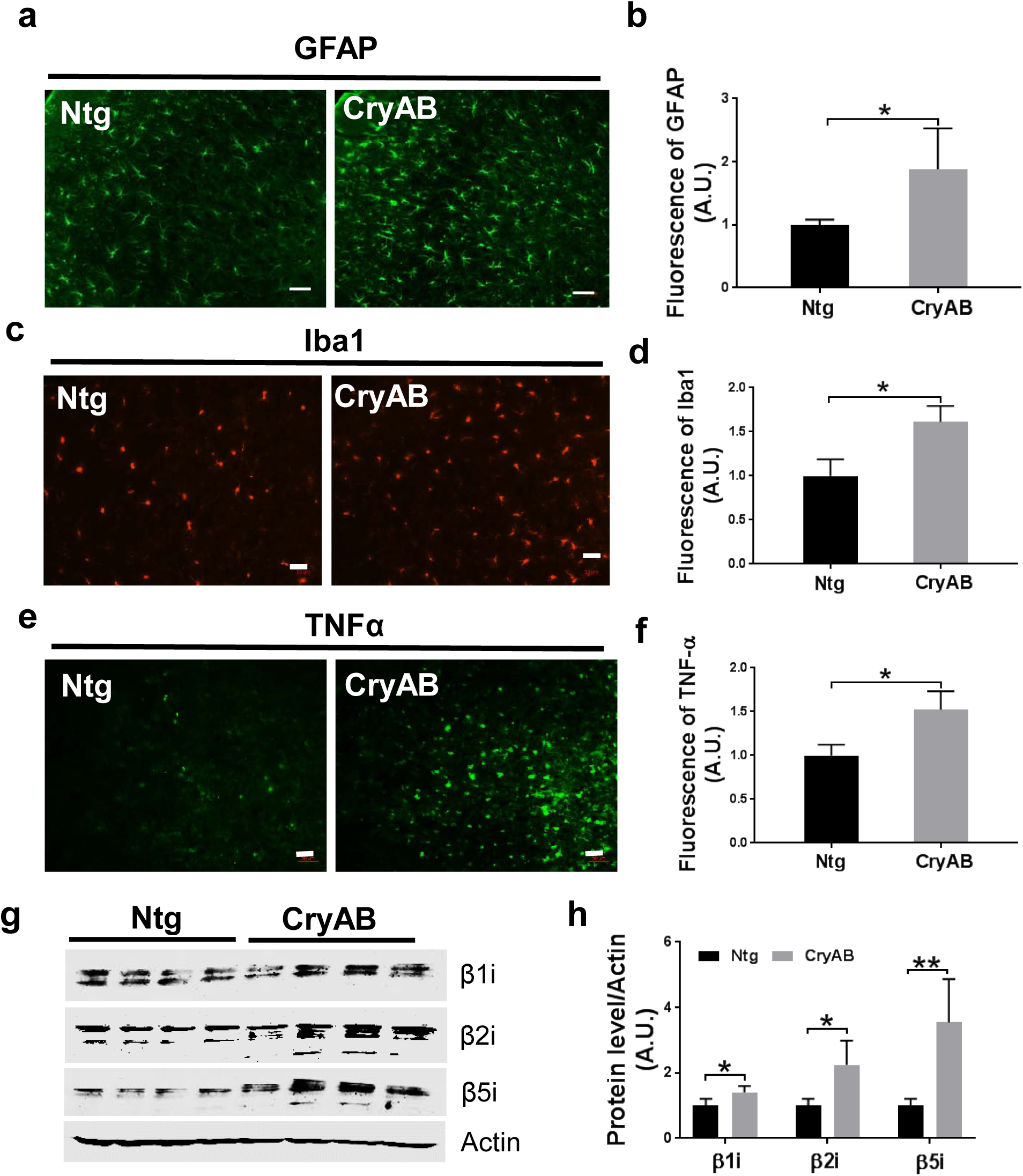
Cardiomyocyte-restricted expression of misfolded CryAB^R120G^ protein enhanced I/R-induced neuroinflammation in the brain. **a** Immunofluorescence staining of GFAP in the brain of mice after MCAO. **b** Quantitative analysis of **a**. **c** Immunofluorescence staining of Iba1 in the brain of mice after MCAO. **d** Quantitative analysis of **c**. **e** Immunofluorescence of TNFα in the brain of mice after MCAO. **f** Quantitative analysis of **e**. **g** Western blot analysis of immunoproteasomes β1i, β2i, and β5i levels in the brain of mice after MCAO. **h** Quantitative analysis of **g**. Data are shown as mean ± SD; n = 3 or 4. * *p* < 0.05, ** *p* < 0.01.

### Cardiomyocyte-restricted expression of misfolded CryAB^R120G^ protein enhanced I/R induced protein aggregation in the brain

As I/R impairs the proteasome and causes protein aggregation in the brain^7–9^, we next determined whether cardiomyocyte-restricted expression of misfolded CryAB^R120G^ influences I/R induced protein aggregation in the brain. Accordingly, we performed Thioflavin-S staining to the Ntg and CryAB^R120G^ mouse brains following 7 days of I/R and observed brighter fluorescence in the CryAB^R120G^ mouse brain than in the Ntg brain (Fig. 5a, b). Notably, Thioflavin-S fluorescence intensity in the I/R induced mouse brains is dramatically brighter than the non-stroke mouse brains (compare Fig. 2a, b with Fig. 5a, b). We then isolated the Triton-X100 insoluble aggregates from the Ntg and CryAB^R120G^ brains following I/R and examined their morphology under an electron microscope. In support of Thioflavin-S staining results, protein aggregates isolated from the CryAB^R120G^ mouse brains were larger and showed more cluster-like features than those isolated from Ntg mouse brains (Fig. 5c). Western blot analysis of the protein aggregates following sonication in an SDS-containing buffer demonstrated that the proteins were (poly)ubiquitinated, with a higher level of (poly)ubiquitin in CryAB^R120G^ mouse brain than in Ntg brain (Fig. 5d, e). Immunohistochemical staining confirmed the colocalization of CryAB protein with (poly)ubiquitin protein in CryAB^R120G^ mouse brains, and both CryAB and (poly)ubiquitin fluorescence intensities are higher in CryAB^R120G^ mouse brains than in Ntg mouse brains at day 7 following I/R (Fig. 5f, g). NeuN immunoreactivity in CryAB^R120G^ mouse brains was reduced at day 7 after I/R (Fig. 5h, i), indicative of neuronal loss, while GFAP and Iba1 immunoreactivities were dramatically increased (Fig. 5j-m), suggesting activation of astrocytes and microglia. Moreover, both GFAP and Iba1 showed a relatively high degree of colocalization with CryAB aggregates, indicating that at this stage after I/R, CryAB aggregates are mainly in astrocytes and microglia. Therefore, cardiomyocyte-restricted expression of misfolded CryAB^R120G^ protein facilitates I/R induced protein aggregation in the brain.

**Fig. 5.**
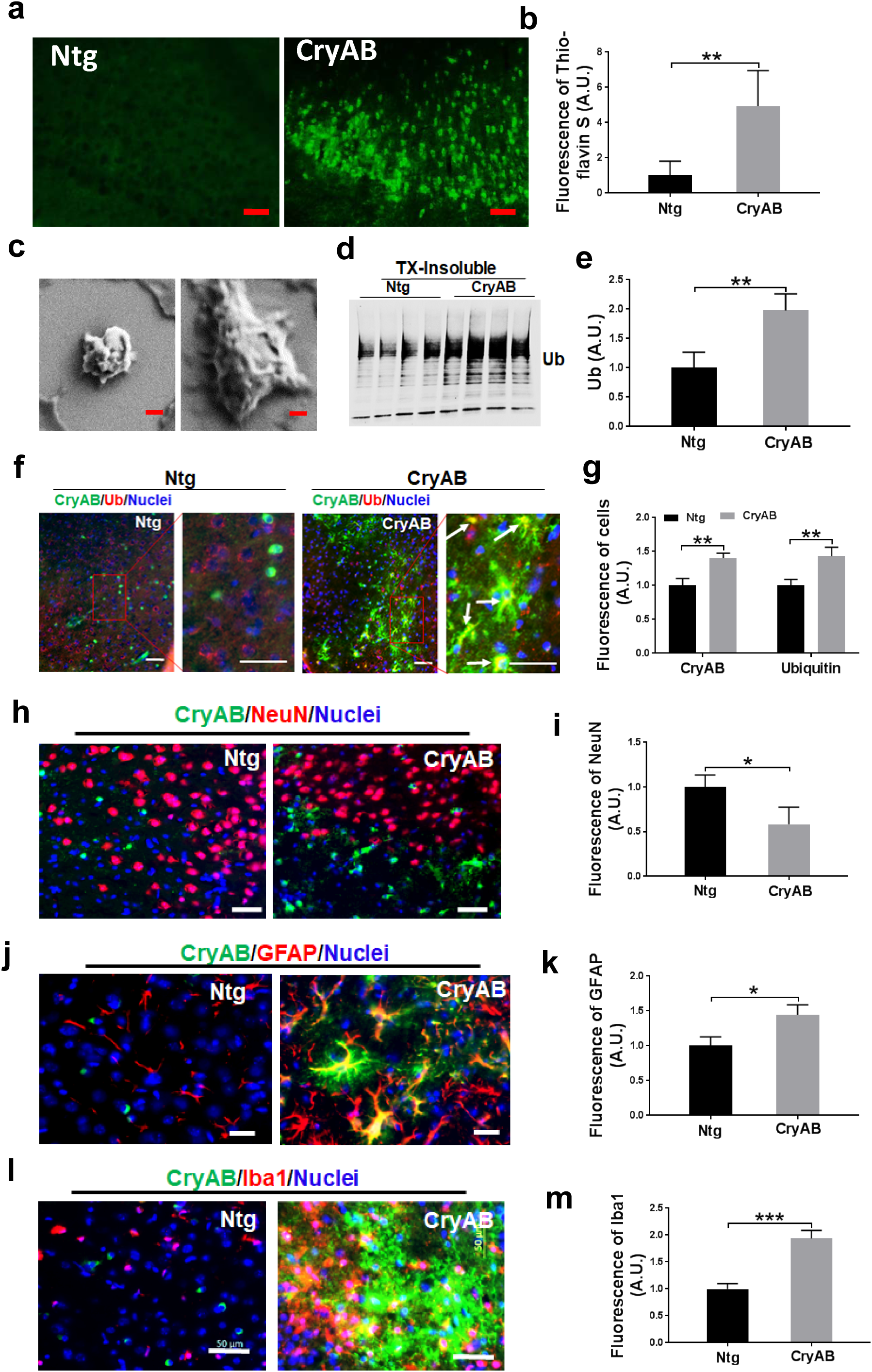
Cardiomyocyte-restricted expression of misfolded CryAB^R120G^ protein enhanced I/R-induced protein aggregation in the brain. **a** Thioflavin S staining of mouse brain at day 7 following MCAO. Scale bar, 50 µm. **b** Quantitative analysis of **a**. **c** Insoluble protein aggregates from mouse brains were analyzed with a scanning electron microscope. Scale bar, 400 nm. **d** Western blot analysis of ubiquitin (Ub) protein level from the Triton X100-insoluble CryAB proteins in the brain of mice after MCAO. **e** Quantitative analysis of **d**. **f** Co-localization of (poly)ubiquitin and CryAB in the brain of mice after MCAO. **g** Quantitative analysis of **f**. **h** Co-staining of NeuN and CryAB in the brain of mice after MCAO. **i** Quantitative analysis of **h**. **j** Co-staining of GFAP and CryAB in the brain of mice after MCAO. **k** Quantitative analysis of **j**. **l** Co-staining of Iba1and CryAB in the brain of mice after MCAO. **m** Quantitative analysis of **l**. Numerical data are shown as mean ± SD; n = 3 or 4. * *p* < 0.05, ** *p* < 0.01, *** p <0.001.

### CryAB^R120G^ mice-derived exosomes disrupt mouse motor and cognitive function following administration to *wild-type* (WT) mice

As the exosome, one type of extracellular vesicles that contain numerous proteins, lipids, RNAs, and other substances, plays a crucial role in mediating cell-cell and heart-brain communications^26, 27^, we next determined whether the increased protein aggregates seen in the CryAB^R120G^ mice were mediated by the circulating exosomes. We isolated exosomes from the blood of both CryAB^R120G^ mice and their Ntg littermates and initially characterized them. Exosomes isolated from both CryAB^R120G^ mouse blood (hereafter referred to as CryAB^R120G^ exosomes) and those from Ntg mouse blood (referred to as Ntg exosomes) expressed exosome-markers, such as Argo1 and Tsg101, despite a remarkably higher level of these marker proteins in CryAB^R120G^ exosomes than in Ntg exosomes (Fig. 6a). Intriguingly, the level of CryAB protein was also dramatically higher in CryAB^R120G^ exosomes than in Ntg exosomes (Fig. 6a). However, the concentration (Fig. 6b, c) and size (Fig. 6b, d) of exosomes in the two types of mouse blood did not differ.

**Fig. 6.**
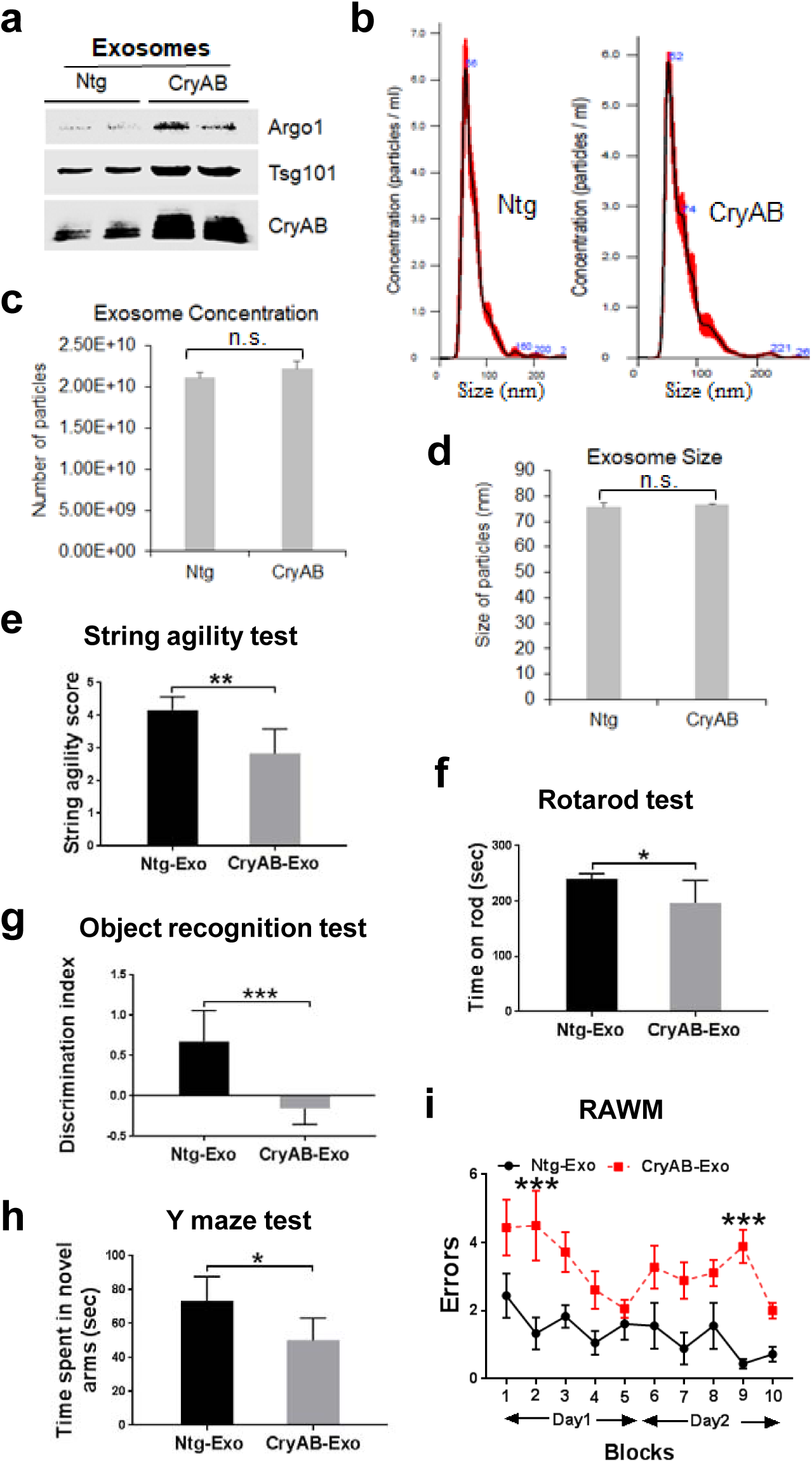
Treatment of WT mice with CryAB^R120G^ mice-derived exosomes disrupted brain functions. **a** The exosomes isolated from the blood of CryAB^R120G^mice expressed the exosome marker proteins such as Argonaut 1 (Argo1) and Tsg101. **b** NTA analysis of exosomes isolated from Ntg (left panel) or CryAB^R120G^ (right panel) mice**. c** The concentration of the CryAB^R120G^ and Ntg exosomes did not differ each other. **d** The size of the CryAB^R120G^ and Ntg exosomes did not differ each other. **e** String agility test result showing impaired animals’ motor function in the WT mice intravenously injected with CryAB^R120G^ derived exosome (CryAB-Exo) compared with the WT mice injected with Ntg-Exo. **f** Rotarod test result showing impaired animals’ motor function in the WT mice intravenously injected with CryAB-Exo compared with the WT mice injected with Ntg-Exo. **g** Objective recognition test result showing reduced memory in the WT mice intravenously injected with CryAB-Exo compared with the WT mice injected with Ntg-Exo. **h** Y maze test result showing reduced memory in the WT mice intravenously injected with CryAB-Exo compared with the WT mice injected with Ntg-Exo. **i** RAWM test result showing reduced memory in the WT mice intravenously injected with CryAB-Exo compared with the WT mice injected with Ntg-Exo. Data are shown as mean ± SD; n = 3 for **a-d** and n = 6 for **e-I.** * *p* < 0.05, ** *p* < 0.01, *** *p* < 0.001.

To test the effects of these isolated exosomes on mice, we intravenously injected the exosomes into the WT C57BL/6J mice (at 2-3 m) daily and performed behavior tests after 2-3 weeks following the treatment. Our results showed that the mice intravenously injected with CryAB^R120G^ exosomes (CryAB-Exo) exhibited reduced motor function, as evidenced by the string agility test (Fig. 6e) and rotarod test (Fig. 6f) compared with the WT animals intravenously injected with Ntg exosomes (Ntg-Exo). Furthermore, results from the novel object recognition test (Fig. 6g), Y-maze test (Fig. 6h) as well as the RAWM test (Fig. 6i) indicated that the WT mice intravenously injected with CryAB-Exo also showed impaired cognitive function in relative to the mice intravenously injected with Ntg-Exo. Thus, CryAB^R120G^ mice-derived exosomes are sufficient to induce motor and cognitive dysfunction in WT mice, supporting the notion that peripherally expressed misfolded proteins can impair brain function through the exosomes.

### CryAB^R120G^ mice-derived exosomes exacerbated I/R-induced brain injury and neurological deficits after administered WT mice

With the exosomes, we then examined whether differential exosomes showed distinct effects on I/R induced brain injury and neurological deficits following I/R in WT mice. Toward this end, WT mice were subjected to 1 h MCAO and after 1 h of reperfusion, the mice were intravenously treated with CryAB^R120G^ exosomes or Ntg exosomes before they were sacrificed at 24 h to assess neuronal injury or were allowed to survive for 7 days to evaluate their survival and functional recovery (Fig. 7a). The mice injected with CryAB^R120G^ exosomes showed increased infarct volume (Fig. 7b, c), reduced survival rate (Fig. 7d), and increased neurological deficits (Fig. 7e) when compared with those injected with Ntg exosomes. Hence, these data support that the exosomes, at least partially, mediate the transfer of the mutant CryAB^R120G^ proteotoxicity from the heart to brain, leading to increased brain injury and neurological deficits.

**Fig. 7.**
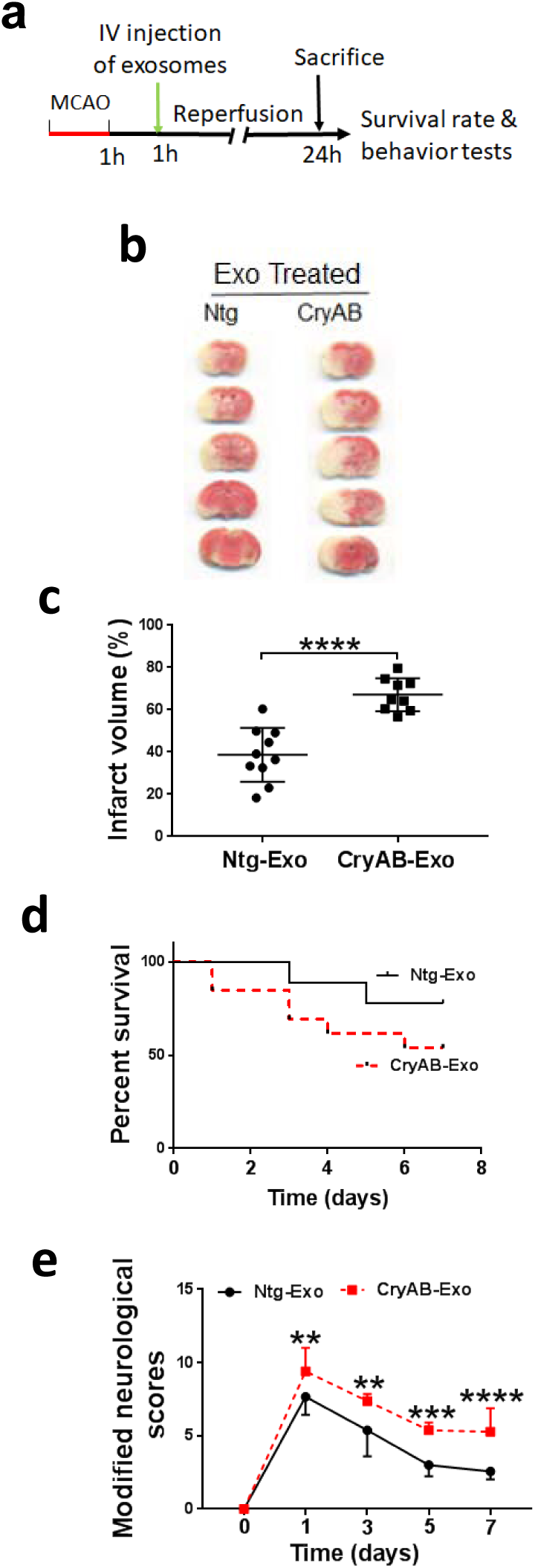
Treatment of WT mice with CryAB^R120G^ mice-derived exosomes worsened I/R-induced brain injury. **a** Diagrammatic illustration of the experimental design of treatment of the wild-type of mice with exosomes. **b** TTC staining of WT mouse brains treated with exosomes isolated from Ntg mice or CryAB^R120G^ mice. **c** Quantitative analysis of **b**. **d** Survival rate of mice following MCAO and exosome treatment. **e** Neurological deficits following MCAO and exosome treatment. Data are shown as mean ± SD; n = 10 for Ntg-Exo treatment and n = 9 for CryAB-Exo treatment in **c-e**. ** *p* < 0.01, *** *p* < 0.001, **** *p* < 0.0001.

### The misfolded CryAB^R120G^ protein showed prion-like propagation when transduced into cultured cells

To further understand how the misfolded CryAB^R120G^ protein induces neuronal injury in the brain, we then treated the primary mouse cortical neuronal cultures with the two types of exosomes. After 24 h following the incubation of exosomes with cultured cells, pronounced protein aggregation were observed selectively in the cells incubated with CryAB^R120G^ exosomes but not with Ntg exosomes (Fig. 8a, b). CryAB^R120G^ exosome-induced protein aggregates were also associated with reduced-ATP level in the neurons (Fig. 8c), suggesting possible mitochondrial dysfunction occurred in the cells.

**Fig. 8.**
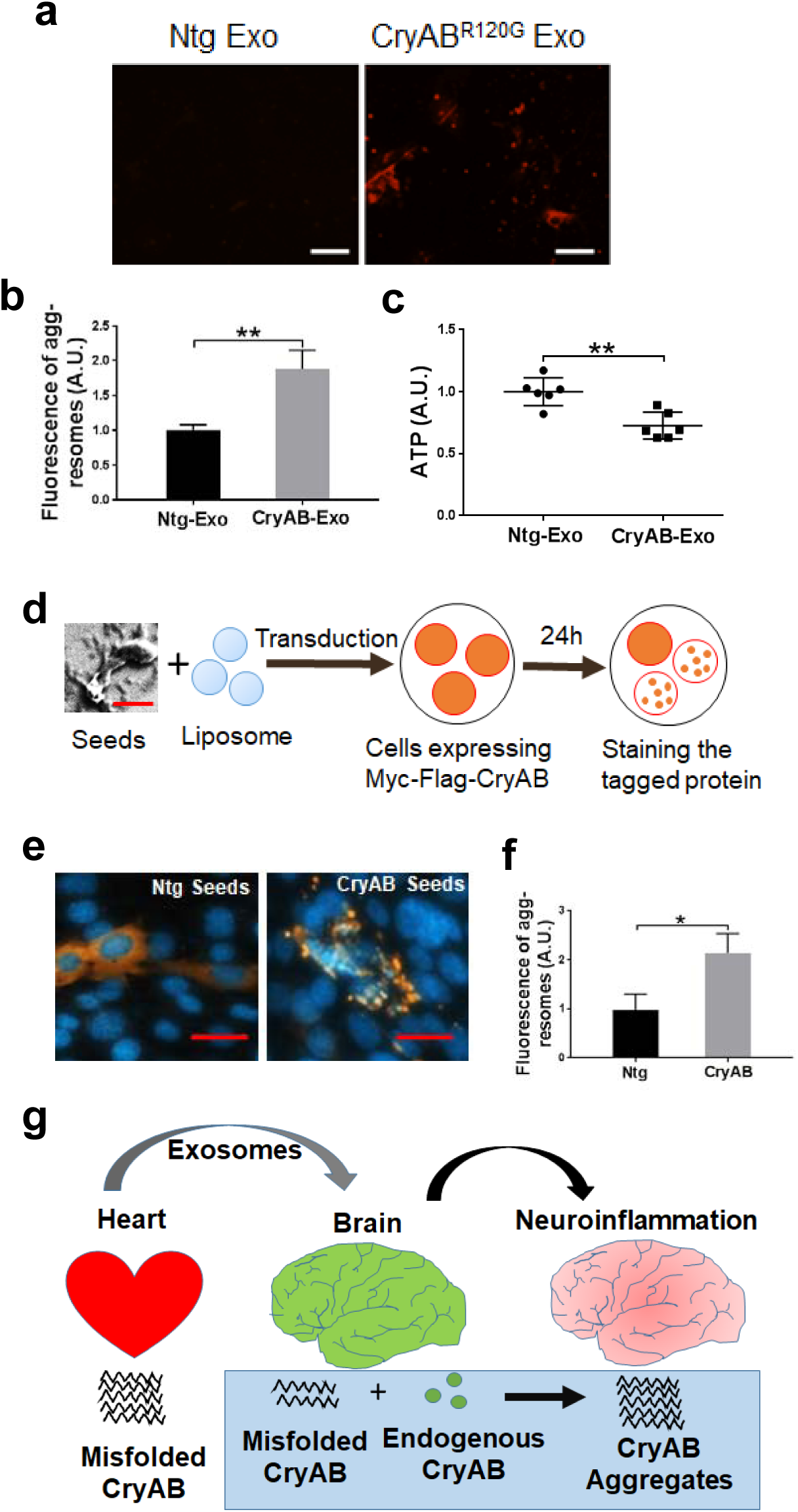
The Prion-like propagation of CryAB^R120G^ protein in cell cultures. **a** Protein aggregates in primary neuronal cultures detected with a Proteostat aggresome detection kit following incubation with the exosomes isolated from CryAB^R120G^ mice. **b** Quantitative analysis of **a**. **c** Cultured neurons treated with CryAB^R120G^ exosomes showed a reduced ATP level. **d** Schematic illustration of an cell culture assay to determine the prion-like property of CryAB^R120G^ protein. **e** Immunocytochemical staining of transfected myc-flag-tagged wild-type mouse CyAB in neural cells following transduction of CryAB^R120G^ aggregate seeds. **f** Quantitative analysis of **e**. **g** Hypothetical model to explain the prion-like propagation of misfolded CryAB^R120G^ protein from the heart to brain. Exosomes containing CryAB^R120G^ protein can be transported from the heart to brain through blood circulation. It is also possible that CryAB^R120G^ protein disrupts BBB and causes CryAB^R120G^ crossing the BBB and entering the brain. In the brain, CryAB^R120G^ protein propagates in a prion-like phenomenon, leading to disruption of BBB, increased neuroinflammation and abnormal brain function. Data are shown as mean ± SD; n = 6 for **c** and n = 3 for **b, f**. * *p* < 0.05, *\*\* p* < 0.01.

As mutant CryAB^R120G^ proteins show dramatic aggregation and form large inclusions in the heart^20^, we finally determined whether the misfolded CryAB^R120G^ translocated from the heart to brain showed the prion-like propagation property in cell cultures. We therefore isolated CryAB^R120G^ protein aggregates from either CryAB^R120G^ mouse or Ntg mouse brains, transduced them into cultured neural cells expressing the *myc*-tagged WT mouse CryAB and then the cells were stained for analyzing the *myc*-tagged CryAB protein aggregation status 24 h following the transduction (Fig. 8d). Transduction of the aggregate seeds into cells was performed by mixing isolated CryAB^R120G^ protein seeds with liposomes before treating cells, as previous data have shown that this would facilitate direct transduction of seeds into the cultured cells, thereby maximizing seed detection efficiency^28^. As expected, the cells receiving CryAB^R120G^ aggregate seeds exhibited remarkable CryAB aggregates (Fig. 8e, right panel), while those receiving Ntg protein aggregate seeds showed diffused CryAB staining (Fig. 8e, left panel). Moreover, the fluorescence intensity in the cell transduced with CryAB protein aggregate seeds was significantly higher than those transduced with Ntg seeds (Fig. 8f). Taken together, these data strongly suggest that the mutant CryAB^R120G^ protein is able to translocate to the brain from the heart via exosomes or directly disrupting BBB function, where it induces prion-like propagation and thereby results in formation of large CryAB aggregates, disruption of neuronal survival and function, and increased neuroinfammation (Fig. 8g).

## Discussion

We here demonstrated that without any insults, cardiomyocyte-restricted expression of the misfolded protein, CryAB^R120G^, is sufficient to impair mouse cognitive and motor function and cause a series of pathological alterations, including formation of protein aggregates, neuroinflammation, BBB dysfunction, and impaired proteasome in the brain. Following acute I/R-induced brain injury, CryAB^R120G^ mice showed increased infarct volume and neurological deficits, enhanced neuroinflammation, and formation of large CryAB protein aggregates positive in ubiquitin staining. Moreover, we further demonstrated that WT mice intravenously injected with the exosomes isolated from CryAB^R120G^ mouse blood showed the similar phenotypes as CryAB^R120G^ mice. Addition of either the purified CryAB^R120G^ exosomes or protein aggregates into cell cultures induced pronounced protein aggregates, exhibiting the prion-like pathological propagation of the misfolded protein. Therefore, our data highlight the significance of peripheral proteostasis in influencing brain functions.

It has been recognized that the communication between the heart and brain is a dynamic and two-way dialogue, with each organ continuously influencing the other’s function^29^. One phenomenon we demonstrated here was that the CryAB^R120G^ mice showed impaired cognitive and motor functions even at 3 m of age, when their cardiac function was at the compensatory stage^20^. Given the fact that the animal’s cardiac contractile function was not reduced at this stage^20^, the observed neurological deficits should not be caused by insufficient cerebral blood supply. Our data support that misfolded CryAB^R120G^ proteins derived from the heart contribute, at least partially, to the impaired brain functions. First, we observed BBB dysfunction in CryAB^R120G^ mice. In mouse model studies, impaired BBB has been found to play a key role in the pathogenesis of numerous neurodegenerative diseases, including Huntington’s disease^30^ and Alzheimer’s disease^31^. Impaired BBB will lead to influx of peripheral macrophages and neutrophils and blood-derived other substances, causing oxidative stress and neuroinflammation^32^. Indeed, we observed increased activation of microglia and upregulation of immunoproteasomes in CryAB^R120G^ mouse brains at 5 m of age in absence of any additional insult. Second, WT mice developed the similar phenotypes as CryAB^R120G^ mice after they were peripherally treated with CryAB^R120G^ exosomes that contained a high level of CryAB protein, presumably the CryAB^R120G^ protein. Finally, CryAB^R120G^ exacerbated I/R-inducted brain injury and neurological deficits. Hence, our results strongly support the role of heart-to-brain translocation of CryAB^R120G^ proteins in triggering brain dysfunction and pathological alterations observed here.

The misfolded CryAB^R120G^ protein enters the brain likely using two pathways, by exosomes and through disrupting BBB. There is compelling evidence supporting that exosomes released by cells in the periphery are able to pass the blood-brain barrier (BBB) and deliver the cargos into brain cells, in particular under inflammatory conditions^33^. Alternatively, the mutant CryAB protein may first enter endothelia of the brain-blood vessels to disrupt their proteostasis and impair the BBB integrity before directly crossing BBB. Once inside brain cells, mutant CryAB^R120G^ proteins undergo the prion-like propagation of protein misfolding, recruiting normal CryAB protein in brain cells to induce CryAB aggregation (Fig. 8g). Although the astrocytes of mouse brains contain more CryAB than neurons^34^, we found that increased CryAB immunoactivity was predominantly located in both microglia and neurons in CryAB^R120G^ mouse brains but not in astrocytes (data not shown) in absence of any insults. Following I/R, however, remarkably increased CryAB immunoactivity was seen mainly in astrocytes and microglia but not in neurons. This phenomenon may result from selective loss of the neurons with the uptake of CryAB^R120G^ protein, due to the nature of its toxicity, especially in the context of an I/R condition. Thus, the heart-derived CryAB^R120G^ proteins induce neuronal damage and glial activation in a prion-like dependent manner, and following neuronal death, the debris including CryAB^R120G^ aggregates were further ingested by astrocytes and microglia to induce activation of additional glial cells, causing neuroinflammation and brain dysfunction (Fig. 8g).

Following I/R, we observed a greater impact of CryAB^R120G^ proteins on brain dysfunction and pathological alterations, including increased infarct size and neurological deficits as well as pronounced neuroinflammation and dramatic accumulation of large protein aggregates. This exacerbation of I/R induced brain injury by cardiomyocyte derived CryAB^R120G^ proteins was likely due to enhancement of the prion-like propagation of CryAB^R120G^ misfolding in the oxidative stress condition. When the isolated CryAB^R120G^ exosomes were added into primary neuronal cultures, they induced formation of large protein aggregates. After transduced into cell cultures, the CryAB^R120G^ mouse brain aggregate seeds triggered normal CryAB proteins to become misfolded and form large protein aggregates. Therefore, these results strongly suggest that a peripherally misfolded proteins disrupt brain function and worsen I/R-induced brain injury through the prion-like pathological mechanism.

In conclusion, we have here demonstrated that without any insult, a peripherally misfolded protein in the heart can impair the animal’s cognitive and motor functions and trigger a series of pathological alterations, including formation of protein aggregates, neuroinflammation, reduced proteasome activity, and BBB dysfunction in the brain. Following I/R, the misfolded CryAB^R120G^ proteins can further worsen brain dysfunction and pathological alterations in the mice. Furthermore, we also have demonstrated the role the exosomes in mediating CryAB^R120G^ proteotoxicity in both the mice and primary neuronal culture.

Importantly, we provided strong evidence that the misfolded CryAB^R120G^ protein aggregate seeds are able to induce the normal CryAB protein into misfolded protein to form large protein aggregates in the prion-like propagation of the proteotoxicity in cell culture. These results suggest that the peripherally expressed misfolded proteins can disrupt brain function and exacerbate I/R induced brain injury through prion-like dependent neuropathology, which may represent an underappreciated mechanism underlying heart-brain crosstalk.

## Methods

### Mice

Adult C57BL/6J wild-type (WT) mice (8–12 weeks of age; mean body weight, 25 g) were purchased from the Jackson Laboratories. CryAB^R120G^ mice and their non-transgenic (Ntg) littermate mice have been previously described^20, 35^. All mice were housed in 3-4 per cage on 12 h light/dark cycle and were given free access to food and water. All animal procedures were approved by the Institutional Animal Care and Use Committee at the University of South Dakota and were in accordance with the National Institute of Health Guide for the Care and Use of Laboratory Animals.

### Transient middle cerebral artery occlusion (MCAO)

The CryAB^R120G^, Ntg, or WT mice animals (male, at 2-3 m) were anesthetized with isoflurane and subjected either to MCAO or sham operation. Transient MCAO was induced by transient occlusion of the MCA as previously described^9, 36^ using a modified intraluminal filament (Doccol, #7020910PK5Re). In brief, male mice were anesthetized with 2% isoflurane and body temperature was maintained at 37.0 ± 0.5°C with an electronic thermostat-controlled warming blanket (Stoelting). Following a permanent of ligation of the external carotid artery (ECA), the filament was inserted into the ECA and then guided toward to the internal carotid artery through the common carotid artery to block the MCA. After 45 min (for the CryAB^R120G^ and Ntg mice) or 1 h (for C57BL/6J WT mice) of MCAO, the occluding filament was withdrawn to allow blood reperfusion. The blood flow of mice were monitored by a Doppler blood flowmeter (Vasamed). After animals were returned to their home cages, they were monitored closely over the next 4 h and then daily for the rest of the study.

### Measurement of infarct volumes

At 24 hours following I/R, mice brains were rapidly removed and stored at -20 °C for 15 minutes to slightly harden the tissue. The brains were sliced into 2-mm thick coronal sections and then incubated with 2% 2,3,5-triphenyltetrazolium chloride (TTC, Sigma, #T8877) to evaluate the infarct volume, as described previously^37, 38^. The infarct volume was manually analyzed using the ImageJ software (NIH).

### Assessment of neurological deficit scores after MCAO

Functional recovery test was performed at 1, 3, 5 and 7 days following MCAO, using a modified neurological severity score (mNSS) system that contains a battery of motor, sensory, reflex, and balance tests.^39^ The mNSS was graded on a scale of 0 to 18 (normal neurological behavior score, 0; maximal neurological deficit, 18): 13-18, severe injury; 7-12, moderate injury; 1-6 mild injury.^39^

### Radial arm water maze (RAWM)

RAWM was performed to assess animals’ learning and memory capability according to the previously described protocol^40, 41^. Briefly, each mouse was gently placed into a swim arm and allowed to find the platform located in the goal arm. On day 1, mice were trained for 15 trials, visible and hidden platform up to 12 trials and the final 3 trials were hidden. On day 2, all 15 trials with hidden platform. The number of incorrect arm entries (errors) was counted until mice found the goal arm or up to 60 seconds. The experiment performer was blind to the genotypes of mice at the time of testing.

### Y-maze analysis

Y-maze analysis was performed to evaluate learning and memory as described previously^40^. Briefly, each mouse is allowed to move freely in start and other arms of Y-maze for 5 minutes (training phase). After 1 hour of rest period, mouse was placed in start arm and allowed to explore all three arms (testing phase). Then, numbers of entries to each arms were recorded for 2 minutes. An entry into an arm was considered when mouse placed all the four paws/legs inside an arm.

### Object recognition test

Mice were tested in a square wooden box (50 × 50 × 30 cm) and habituated to the empty box for 5 minutes the day before the test. On the first day of test, mice were presented with two similar objects in the box and permitted to explore freely for 10 minutes. On the second day of test, one of the two familiar objects was randomly replaced by a new one and mice were placed back into the box and allowed to explore the objects for 10 minutes. The amount of time spent to explore each object was recorded, and the relative exploration of the novel object was presented by a discrimination index (DI = (T_novel_ - T_familiar_)/(T_novel_ + T_familar_).

### Open field test

Open field test was used to assess mouse activity and exploratory behavior. Briefly, each mouse was placed to the center of an open box (81 × 81 × 28.5 cm), and the box floor contained four horizontal and four vertical lines to demarcate 16 squares (20 × 20 cm each) and mice were allowed to explore the interior for 5 min. The total number of line crossings and time spent in the center and peripheral areas were recorded.

### String agility test

This test was used to test animals’ agility and grip capacity^40^. Briefly, each mouse was allowed to grasp a suspended string only using their forepaws and subsequently released by the experimenter. The maximum trial length was 60 seconds. The scoring system to assess the grasp capacity involved a five-scale system: 0-unable to remain on string; 1-hangs by two forepaws; 2-attempts to climb to string; 3-hangs by two forepaws and one or two hind paws around string; 4-four paws and tail around string; 5-escape.

### Rotarod test

This was used to evaluate mouse balance and general motor function and was conducted using an accelerating Rotarod machine^40^ (Med Associates). Each animal was placed on an accelerating Rotarod and trained for three successive days with 3 trials each day. The maximum trial length was 250 seconds. The 4th day was tested with 3 trials for each mouse.

### Thioflavin S staining

Thioflavin S staining of protein aggregates in the brain was performed using a previously described method^40^. Brain sections were incubated with 70% ethanol (Fisher, #A40520) for 1 minute and then 80% ethanol for 1 min. Slides were then incubated with 0.1% Thioflavin S (Sigma, #T1892) in 80% ethanol for 15 minutes. Slides were then rinsed with 80% ethanol for 1 minute, 70% ethanol for 1 minute and distilled water two times. Slides were mounted with Cytoseal 60 Medium (Richard-Allan Scientific, #8310-16) and images were acquired using a fluorescence microscope equipped with the ZEN 2.5 software (Carl Zeiss, Jena, Germany).

### Nissl staining

Brain sections that were rehydrated through 100% and 95 % alcohol to distilled water were stained in 0.1% Cresyl Violet (Acros Organics, #10510-54-0) solution for 5-10 minutes and then rinsed quickly in distilled water before were differentiated in 95% ethanol, for 20 minutes and dehydrated in 100% ethanol 2 times, each for 5 min. Finally, the sections were cleared in Xylene for 5 min twice and were mounted with Cytoseal 60 Medium (Richard-Allan Scientific, #8310-16). Images were captured under a microscope (Olympus). The number of surviving neurons was quantified by using the ImageJ software (NIH).

### Immunofluorescent staining

At 2 or 7 days post-MCAO, the PBS-perfused and paraformaldehyde-fixed brains were coronally sliced into 10 μm thickness sections on a Cryostat (Leica, Buffalo Grove, IL, USA) at -20 °C. Sections were blocked with 5% bovine serum albumin (Fisher, # BP9706-100) and then incubated with anti-GFAP (1:1000, EMD Millipore, #AB5804; Lot 2013929), anti-Iba1 antibody (1:1000, FUJIFILM Wako, #019-19741; Lot LKF6437), anti-NeuN antibody (1:1000, Millipore, #ABN78, Lot 1979271), anti-CryAB antibody (1:100, Abcam, # ab13496), anti-TNF alpha antibody (1:100, Abcam, # ab1793, Lot GR3261538-1), anti-Myc antibody (Cell Signaling Technology, # 2276, Lot 24) or anti-ubiquitin antibody (1:500, Abcam, # ab7780, Lot GR3196708-1). This was followed by incubation with Cy3- or Cy2-conjugated goat anti-rabbit or goat anti-mouse secondary antibodies (Jackson ImmunoResearch, #711-165-152) or Dylight 488 conjugated IgG, #35552, Lot QK224422). In some experiments, nuclei were counter-stained with a DNA binding dye, Hoechst 33342 (1:1000, ThermoFisher, #H3570). Images were captured with a fluorescence microscope equipped with the ZEN 2.5 software (Carl Zeiss).

### Analysis of BBB permeability

CryAB and Ntg mice were intraperitoneally injected with 800 µl of 1% (w/v) Evans blue dye (Sigma, #E2129). After 1 h following the injection, the mouse blood was collected and the animals were perfused with a phosphate-buffered saline (PBS) before the brains were collected. Both the blood and the brains were used for evaluation of BBB permeability as described previously.^42^ Briefly, brains were cut into small pieces (±50 mg) and incubated with 0.9% saline and then homogenized in 1:3 volume-ratios of 50% trichloroacetic acid (TCA, Sigma, #T9159). The blood samples were centrifuged at 2500 × g for 15 min to obtain plasma samples and the treated with 1:3 volume-ratios of 50% TCA. Subsequently, the samples were further centrifuged at 10,000 × g for 20 min. The supernatants (30 µl) and 90 µl of 95% ethanol were add to each well of 96 well plate and the dye was quantified at 460 nm excitation/535 nm emission on a microplate reader (PerkinElmer) for the brain samples or measured at 620 nm excitation/680 nm emission with a BioTek plate reader (BioTek Instruments) for the blood samples.

### Western blot

Electrophoresis, transfer and western blot analysis of proteins were according to previously described methods^43^. The primary antibodies used were anti-CryAB (1:1000, Abcam; #ab13496), anti-proteasome 20S beta 1i subunit (1:1000, Enzo life sciences, #BML-PW8205; Lot 10071464), anti-proteasome 20S beta 2i subunit (1:1000, Enzo life sciences, #BML-PW8350; Lot 11281405), anti-proteasome 20S beta 5i subunit (1:1000, Cell Signaling Technology, #13726; Lot 1), anti-Argonaute 1 (Argo1) (1:1000, Cell Signaling Technology, #5053, Lot 1), anti-Tsg101 (1:1000, Santa Cruz Biotechnology, #sc7964, Lot F0217), anti-ubiquitin (Ub) (1:1000, Cell Signaling Technology, #3936, Lot 14), anti-K48-linkage specific polyubiquitin (1:1000, Cell Signaling Technology, #8081; Lot 2), and anti-actin (1:1000, Santa Cruz Biotechnology, #sc-1616, Lot G1615). The secondary antibodies used were conjugated with the infrared dyes (1:5000, LI-COR, #926-32211, Lot C80118-05; #92668072, Lot C71204-03; #926-33214, Lot C80207-07). Protein band intensities were measured using an Odyssey scanner (LI-COR) and quantified using UN-SCAN-IT gel6.1 software (Silk Scientific).

### Isolation of exosomes from mouse blood plasma

Whole blood was collected from facial vein of mice. Isolation of exosomes was performed as described previously with minor modification^44^. Briefly, 0.8 µm-filtered blood plasma samples were diluted by two folds with PBS and then centrifuged at 13,200 × g at 4 °C for 22 min to remove microvesicles. The supernatant was filtered twice through 0.22 µm-filters. Exosomes were pelleted by ultracentrifugation at 120,000 × g with a SW41Ti rotor (Beckman Coulter). Pellets were washed once with PBS and protein concentration was determined by the NanoDrop^TM^ 2000/2000c spectrophotometers (Thermo Fisher).

### Nanoparticle Tracking Analysis (NTA) of exosomes and treating WT mice with the exosomes

Prior to NTA, samples were diluted at 1:100 in PBS and 1 ml exosome solution was used for NTA analysis to assess the size and number of EVs using the NanoSight NS300 system (Malvern Instruments).

To determine whether the isolated exosomes influence brain function in WT mice, WT C57BL/6J mice were intravenously injected with 0.8 mg/kg of exosomes daily and animal motor and cognitive behaviors were examined after 2-3 weeks following the treatment using the behavioral test methods mentioned above. To determine the effect of isolated exosomes on I/R induced brain injury, the exosomes (3.75mg/kg) were injected into WT C57BL/6J male mice through tail vein after 1 h of MCAO and following the treatment, the mice were either euthanized after 24 h to assess their brain infarct volume or allowed to survive for 7 days to evaluate their survival rate and neurological deficit scores using methods described above.

### Isolation of protein aggregates and scanning electron microscopy (SEM)

Isolation of Triton X-100 insoluble protein aggregates from Ntg or CryAB^R120G^ mouse brains were according to previously described methods^10, 45^. Examination of isolated protein aggregates with SEM was performed with the field emission SEM (SIGMA, Zeiss). The samples were prepared by drop casting on Si wafer and sputter coated with Au for the imaging. Secondary electron images were acquired by scanning with 1.5-2.0 kV acceleration voltage.

### Primary neuronal culture, exosome treatment, and ATP measurement

Primary mouse cortical neurons were prepared from wild-type C57BL/6J mice as previously described^43^. The cells were cultured in poly-ornithine-coated 12-well plates for 7 days and then treated with exosomes (10 µg/ml) isolated either from the Ntg or CryAB^R120G^mouse blood and after 24 hours, the exosome-treated neuronal cultures were stained for protein aggregates using a Proteostat Aggresome Detection Kit (Enzo Life Sciences, #ENZ-51035) by following the manufacturer provided protocol. To test the effect of exosomes on cell viability, the treated cells were collected for ATP assay using an ATP Bioluminescence Assay Kit CLS II (Sigma, #11699695001) according to the protocol provided by the manufacturer.

### CryAB^R120G^ seeding assay

To determine the prion-like property of mutant CryAB^R120G^ protein, mouse striatal cells (Coriell) or HEK293 cells (ATCC) were cultured in 12-well plates in the complete medium containing the Dulbecco’s Modified Eagle Medium supplemented with 10% fetal bovine serum and penicillin/streptomycin (Thermo Fisher). Cells were transfected with myc-flag-CryAB plasmid (OriGene, #MR201515) encoding the myc-flag-tagged mouse CryAB protein. After 24 hours, the cells were transduced with 500 ng/ml Triton X-100 insoluble aggregates isolated either from Ntg or CryAB^R120G^ mouse brains using Trans-Hi^TM^ transfection reagent (FormuMax, #F90101TH). The transduced cells were fixed and subjected to immunocytochemical staining with an anti-Myc antibody (see above **Immunofluorescent staining** part) 24 hours following the transduction.

### Statistical analysis

Statistical analyses were conducted using GraphPad Prism version 7.0 statistical software. Differences between two groups were assessed using unpaired t test. Significant differences between more than two groups were analyzed using one-way or two – way ANOVA followed by Tukey’s post hoc test or Sidak’s multiple comparisons test. *P* < 0.05 was regarded as statistically significant.

### Data availability

Data supporting the findings of this manuscript are available from the corresponding author upon reasonable request.

## Acknowledgements

The work was supported by the NIH grants HL072166 and HL131667 (X.W.), and NS088084 (H.W.).

## Author contributions

H.W. conceived the project and interpreted data. Y.L., K.S., A. B., R.S., X.W. and H.W. designed experiments. X.W. provided animals. Y.L., K.S., C.C.H., A.B., and R.S. performed experiments, collected and analyzed data. Y.L., K.S. and H.W. wrote the paper. All authors read and edited the paper and approved submission.

